# Spatial distribution of neuropathology and neuroinflammation elucidate the biomechanics of fluid percussion injury

**DOI:** 10.1101/2020.10.05.325514

**Authors:** Joshua A. Beitchman, Jonathan Lifshitz, Neil G. Harris, Theresa C. Thomas, Audrey D. Lafrenaye, Anders Hånell, C. Edward Dixon, John T. Povlishock, Rachel K. Rowe

## Abstract

Diffuse brain injury is better described as multi-focal, where pathology can be found adjacent to seemingly uninjured neural tissue. In experimental diffuse brain injury, pathology and pathophysiology have been reported far more lateral than predicted by the impact site. Finite element biomechanical models of diffuse brain injury predict regions of maximum stress and strain. However, the application of a skull with uniform thickness may mask the pathophysiology due to varying thickness of human and animal skulls. Force applied to the intact skull would diffuse the forces, whereas forces applied through an open skull are distributed along paths of least resistance within and then exiting the skull. We hypothesized that the local thickening of the rodent skull at the temporal ridges serves to focus the intracranial mechanical forces experienced during brain injury and generate predictable pathology in underlying cortical tissue. We demonstrated local thickening of the skull at the temporal ridges using contour analysis of coronal skull sections and oblique sectioning on MRI. After diffuse brain injury induced by midline fluid percussion injury (mFPI), pathological foci along the anterior-posterior length of cortex under the temporal ridges were evident acutely (1, 2, 7 days) and chronically (28 days) post-injury by deposition of argyophilic reaction product. Area CA3 of the hippocampus and lateral nuclei of the thalamus showed pathological change, suggesting that mechanical forces to or from the temporal ridges shear subcortical regions. A proposed model of mFPI biomechanics suggests that injury force vectors reflect off the skull base and radiate toward the temporal ridge due to the material properties of the skull based on thickness, thereby injuring ventral thalamus, dorsolateral hippocampus, and sensorimotor cortex. Surgically thinning the temporal ridge prior to injury reduced the injury-induced inflammation in sensorimotor cortex. These data build evidence for the temporal ridges of the rodent skull to contribute to the observed pathology, whether by focusing extracranial forces to enter the cranium or intracranial forces to escape the cranium. Pre-clinical investigations can take advantage of the predicted pathology to explore injury mechanisms and treatment efficacy.

**Highlights:**

- The temporal ridge is 75% thicker than the adjacent skull of the rodent
- Experimental diffuse TBI neuropathology occurs beneath the length of the temporal ridge
- Neuropathology encompasses sensorimotor cortex, somatosensory thalamus, and dorsolateral hippocampus
- Proposed mechanism of biomechanical injury forces include the temporal ridge

## Introduction

Clinical and experimental traumatic brain injury (TBI) involve both a primary injury and subsequent disease processes that dismantle, repair, and regenerate circuits in the brain [1, 2]. In response to the initial mechanical injury, adaptive repair and regeneration fail to reconstitute the original neuronal circuits, leaving a mis-wired brain and neurological impairments that decrease quality of life. Clinically, this process is relevant to veterans, athletes, survivors of interpersonal violence, children, and the elderly who experience one or more TBIs [3, 4]. We and others apply experimental models of TBI to study the acute and chronic events associated with this disease and use these models to develop therapeutic interventions.

Various models of experimental TBI have been developed, where fluid percussion injury (FPI) is one of the oldest and best characterized animal models of TBI [1, 5-8]. Specifically, the midline FPI (mFPI) model is capable of producing a diffuse, concussive-like, TBI in rodents, whereas lateral FPI (lFPI) produces a mixed focal and diffuse injury [1, 7, 9-11]. To this end, the primary FPI neuropathology is diffuse axonal injury (DAI), rather than overt cell death. The clinical relevance of these injuries includes a transient suppression of neurological reflexes and acute motor deficits [5, 6, 10, 12-28]. In the more chronic phases of injury, evidence of somatic, cognitive, and affective symptoms emerge. Somatic morbidity includes a sensitivity to facial whisker stimulation, similar to agitation in people [29-34]. Cognitive performance is degraded in short, long, and working memory, using various cognitive testing modalities [10, 32, 35-45]. Affective symptoms are identified by disruption of circulating hormone levels and responses in anxiety tests [10, 46, 47]. Thus, FPI affords investigations into the development and maintenance of cognitive, somatic, and affective morbidities, which parallel clinical impairments. Further investigations provide evidence in the forms of diagnostic tests, therapeutic interventions, and rehabilitation to improve healthcare delivery and quality of life for TBI survivors.

Despite the broad implementation of the FPI model in the neurotrauma field, biomechanical models fail to accurately explain the resultant distribution of pathology [48-55]. In the initial implementation of FPI in rodents, Dixon describes that this technique does “*not attempt to reproduce rapid acceleration-deceleration of the head…. Rather, fluid-percussion brain injury successfully produces graded levels of injury associated with predictable neurologic, physiologic and histologic changes that are comparable to those observed in human brain injury*” [14]. In an approach to observe the mechanical forces of injury on the brain, Dixon *et al*. conducted high-speed cineradiographic studies of the brain inside the skull over the 15 msec of mFPI injury [14]. In this way, the primary injury was first characterized by Dixon et al. as “*intracranial fluid movement … by rapid radial movement within the epidural space …. suggesting that the image of the indentation acutely may have been caused by lateral fluid displacement following the curvature of the skull*” [14]. Thereafter, subsequent publications comment on the resultant pathology. Hicks *et al*., reported that “*it is interesting to note that the primary site of cortical damage is ventrolateral, rather than directly underneath the impact site*” [56]. These authors continue to explain this phenomenon as a consequence of biomechanical forces on selectively vulnerable tissue, based on regional or cellular cytoarchitecture. Over the 30+ years of research using FPI in rodents, curious pathology, similar to the observation by Hicks *et al*., has been reported as focal damage far more lateral than predicted by the location of the applied mechanical forces of injury. Today, the biomechanical mechanism for neuropathology localizing far more lateral than the craniotomy remains an enigma. For the midline variant of FPI, where the mechanical forces are applied over the exposed sagittal sinus, the core pathologies are identified millimeters more lateral from the injury site. Further, the cortical pathology does not necessarily align with cytoarchitectural landmarks, as posited by Hicks *et al* [56]. The curious pathology occurs far lateral from the injury site in other non-focal TBI models, such as impact acceleration [57]. In this communication, we reference the range of pathologies occurring far lateral from the injury site to include, but in no way limited to, blood brain barrier disruption, axotomy, plasma membrane permeability, and cell death. Upon reexamination, we recognized that the curious pathology of diffuse TBI tracked beneath the temporal ridge of the skull, where the muscles of mastication adhere.

In the present communication, we make a case for differential thickness along the rodent skull as a contributing factor to the direction of biomechanical forces of diffuse brain injury, in addition to inherent properties of the tissue. Thus, we hypothesized that a local thickening of the rodent skull at the temporal ridges serves to focus mechanical forces of brain injury and generate predictable pathology in line with the temporal ridges. Using peer-reviewed literature, contour analysis of the skull, MRI imaging, and histopathology, we characterized the temporal ridge of the rodent and the development of consistent and persisting neuropathology surmised to result from maximal tissue strain and stress. The succession and alignment of pathology through the cortex, hippocampus, and thalamus predict possible force vectors through the cranial vault. Finally, we propose a new biomechanical model for midline FPI, in which forces are focused onto the tissue by the temporal ridge and provide evidence that reduction of bone thickness limits the development of neuropathology post-injury.

## Methods

### Compendium of experimental traumatic brain injury publications

A compendium of literature was assembled by using experimental brain injury publications for low-power photomicrographs that include primary sites of pathology as photomicrographs or schematics. Histological and radiological techniques that identify pathology were not restricted to a single approach, rather identifying photomicrographs or schematics primarily by complete or hemispheric coronal section. Relevant manuscripts, figure numbers, and histological evidence are presented in table format. Publications were screened from the personal libraries of the primary authors and sequential paging through printed copies of the *Journal of Neurotrauma* (1993-2010). Relevant images were searched on the world wide web (https://images.google.com) for histological or radiographic coronal images of brain-injured rodents. All figures depicting pathological patterns in low power photomicrograph or drawn schematic were included in the compendium.

### Animals

Animal work was conducted using 8-12-week-old male Sprague-Dawley rats. Rats were pair housed in a normal 12 hour light/dark cycle and fed a standard rodent diet. Food and water were available *ad libitum*. All practices were conducted in accordance with the guidelines established by the internal Institutional Animal Care and Use Committee (IACUC) and the National Institute of Health (NIH) Guidelines for the Care and Use of Laboratory Animals. Studies are reported following the Animal Research: Reporting *In Vivo* Experiments (ARRIVE) guidelines [58]. Randomization of animals was achieved by assigning animals to treatment groups before the initiation of the study to ensure equal distribution across groups. Data collection stopped at pre-determined final endpoints based on days post-injury for each animal. Animals were evaluated daily for 3 days for postoperative recovery by a physical examination and documentation of each animal’s condition. Pre-determined exclusion criteria included post-operative weight loss greater than 15% of pre-surgical weight. No rats were excluded from this study.

### Flesh eating beetles

Skulls from 8-12-week-old naïve rats were prepared by dermestid beetles (*dermestes maculatus*). Rat heads were skinned, hung to dry, and then placed in glass jars, with nesting cotton, covered with a moist cloth to increase humidity, and shielded from light. Within 10-14 days, the carrion was cleaned from the skulls by the beetle larvae. Skulls were further cleaned with bleach water and air-dried. Photographs were taken of complete rat skulls and rat skulls that were cut in the coronal plane with a hack saw (exposed surfaces were marked with permanent marker to increase contrast). Measurements were taken along the circumference of the calvarium using calipers, focusing on the medial-lateral midpoint of the parietal bone and the temporal ridge.

### Magnetic resonance imaging (MRI) of the rat head

A cohort of naïve 8-12-week-old rats was anatomically imaged by MRI to visualize the relationship between the brain, skull, and musculature. All data were acquired on a 7 Tesla spectrometer (Oxford Instruments, Oxford, UK) controlled by a Bruker Biospec console (Bruker Biospin MRI Inc, Billerica, MA, USA). The rat was anesthetized with isoflurane (4% induction, 1.5% maintenance, vaporized in oxygen) positioned in a purpose-build plexiglass cradle using a bite-bar and ear bars. Data were acquired using a ^1^H radiofrequency (RF) volume resonator in transmit-only mode, and a pulse-decoupled receive only surface RF coil placed over the head. Optimal field homogeneity was accomplished by ensuring the head was positioned in the center of the magnet, by automated global shimming, and by shim calculation after obtaining the b0 field map over the brain using the Bruker Paravision macro MAPSHIM. Image acquisition was performed using a two-dimensional, rapid acquisition with relaxation enhancement (RARE) pulse sequence using a 35×35 mm field-of-view encoded in a 128×128 data matrix, with 50 coronal image slices 0.5 mm thick, resulting in a resolution of 234 × 234 × 750 µm. The following imaging parameters were used: 6 sec repetition time, 56 msec echo time, 50 kHz bandwidth, 4 averages per phase-encoding increment, and rare factor 8. Data were Fourier transformed into 16-bit signed integer spatial data, and then regrouped into compressed NIFTI format. Image stacks were evaluated using the Volume Viewer 1.31 plugin on NIH Image. Images were rotated, segmented, and pseudocolored to represent relationships between the brain, skull, and musculature with respect to the temporal ridges of the skull.

### Midline fluid percussion injury

Adult male Sprague-Dawley rats were subjected to midline fluid percussion injury (FPI) consistent with methods described previously [9, 21, 59, 60]. Final animal numbers are indicated in the results section for each study. Briefly, rats were anesthetized with 5% isoflurane in 100% O_2_ and maintained at 2%/100% O_2_ via nose cone. During surgery, body temperature was maintained with a Deltaphase^®^ isothermal heating pad (Braintree Scientific Inc., Braintree, MA). In a head holder assembly (Kopf Instrument, Tujunga, CA), a midline scalp incision exposed the skull. A 4.8-mm circular craniotomy was performed (centered on the sagittal suture midway between bregma and lambda) without disrupting the underlying dura or superior sagittal sinus. An injury cap was fabricated from the female portion of a Luer-Loc needle hub, which was cut, beveled, and scored to fit within the craniotomy. A skull screw was secured in a 1-mm hand-drilled hole into the right frontal bone. The injury hub was affixed over the craniotomy using cyanoacrylate gel and methyl-methacrylate (Hygenic Corp., Akron, OH) was applied around the injury hub and screw. The incision was sutured at the anterior and posterior edges and topical lidocaine ointment was applied. Animals were returned to a warmed holding cage and monitored until ambulatory (approximately 60-90 minutes).

For injury induction, animals were re-anesthetized with 5% isoflurane 60-90 minutes after surgery. The dura was inspected through the injury-hub assembly for debris, which was then filled with normal saline and attached to the male end of the fluid percussion device (Custom Design and Fabrication, Virginia Commonwealth University, Richmond, VA). An injury of moderate severity (2.0-2.1 atm; 5-8 minute righting reflex time) was administered by releasing the pendulum onto the fluid-filled cylinder, as reflexive responses returned. Animals were monitored for the presence of a forearm fencing response and the return of the righting reflex as indicators of injury severity [21]. After injury, the injury hub assembly was removed *en bloc*, integrity of the dura was observed, and the incision was stapled. Brain-injured animals had righting reflex recovery times greater than 5 minutes. After recovery of the righting reflex, animals were placed in a warmed holding cage before being returned to their home cages. Adequate measures were taken to minimize pain or discomfort.

### Amino-cupric silver technique

At 1, 2, 7 and 28 days post-injury (DPI) brain-injured rats (n=3/time point) were overdosed with sodium pentobarbital (200 mg/kg i.p.) and transcardially perfused with 0.9% sodium chloride, followed by a fixative solution containing 4% paraformaldehyde. Following decapitation, the heads were stored in a fixative solution containing 15% sucrose for 24 hours, after which the brains were removed, placed in fresh fixative, and shipped for histological processing to Neuroscience Associates Inc. (Knoxville, TN). The rat brains were embedded into a single gelatin block (Multiblock^®^ Technology; Neuroscience Associates). Individual cryosections containing all the rat brains were mounted and stained with the de Olmos aminocupric silver technique according to proprietary protocols (Neuroscience Associates) to reveal argyrophilic reaction product, which localized to neurons and neuronal processes, counterstained with Neutral Red, and then cover-slipped. The stained sections were analyzed in our laboratory. Every sixth section from the anterior commissure through the substantia nigra was imaged at 1.25x, masked from the background and overlaid on the remaining sections from the same brain. Silver quantification of the S1BF and VPM from this tissue has been previously published [61, 62]. Evidence for pathology along the tissue underneath the temporal ridges was noted in all brain-injured animals.

### Shaved temporal ridge of the skull and immunohistochemistry

Similar to the mFPI surgical procedure describe above, a new cohort of rats (n=9) was prepared for injury induction. In addition to the procedures above, none (n=3), the rat’s anatomic left (n=3), or both (n=3) temporal ridge(s) of the skull were shaved by manual scraping to approximate the thickness of the calvarium, thus leaving both temporal ridges intact, the rat’s anatomical right temporal ridge intact, or a skull devoid of temporal ridges, respectively. Rats were randomly assigned to have the temporal ridges shaved. Rats were then administered a moderate FPI as described above. At 7 DPI, rats were given an overdose of sodium pentobarbital and transcardially perfused with 4% paraformaldehyde after a phosphate buffered saline (PBS) flush. Brains were removed and cryoprotected in 30% sucrose and stored frozen. After freezing, brains were cryosectioned in our laboratory in the coronal plane at 20 μm and thaw mounted onto gelatinized glass slides and stored at −80°C. Prior to cryosectioning, a needle was placed through the ventral left hemisphere to accurately identify hemispheres throughout histological processing. Frozen slides were removed from −80°C, placed in an oven at 60°C for approximately 4 hours and then rinsed 3 times for 5 minutes each in PBS. Next, the slides were incubated in 4% goat serum blocking solution for 1 hour. Slides were incubated with the primary antibody (rabbit anti-ionized calcium binding adaptor molecule 1, IBA-1; 1:1000, Item # 0199-19741, Wako Chemicals, Richmond, VA) and stored at 4°C overnight. Slides were rinsed 3 times in PBS and the secondary antibody (biotinylated horse anti-rabbit; 1:250, Vector Laboratories, Burlingame, CA) was added and slides were incubated at room temperature for 1 hour. Slides were rinsed 3 times in PBS, incubated in 30% H_2_O_2_ for approximately 20 minutes and rinsed again in PBS 3 times. Diaminobenzidine (DAB) with H_2_O_2_ and nickel was then prepared in PBS. 300 µL was then applied to each slide for equal duration, until the chromogen reaction was visualized. Slides were washed in tap water, and washed in 70%, 90%, 100% ethanol, each for 5 minutes. Finally, slides were incubated in citrisolve 2 times for 10 minutes and coverslipped using DPX. The immunostained slides were imaged in our laboratory using an Olympus AX80 Automatic Research microscope with attached DP70 digital camera.

### Statistical analysis

Data were organized using Microsoft Excel® and analyzed using Prism® software (Graphpad Software, Inc, La Jolla, CA). Data points collected bilaterally (e.g. thickness of the skull) were averaged to represent a single animal before comparison. A Student’s two-tailed t-test was used to compare values between groups, with significance defined at p<0.05.

## Results

### Peer-reviewed literature identified TBI pathology in cortex beneath the temporal ridge

Cortical pathology beneath the temporal ridge following experimental TBI, particularly FPI in its many variations, has been identified across multiple laboratories, involving numerous surgeons, over at least a decade. In Table 1, we list 46 publications between the years of 1987 and 2010 with a low-power micrograph or summary schematic of pathology localized under the temporal ridge induced by diffuse or mixed model brain injury. The range of pathological assessments (e.g. histological, metabolic, enzymatic, vascular) provides evidence that FPI-induced neuropathology is multi-dimensional and not limited to a single type of neuropathology.

**Table 1.** Table of compiled peer-reviewed literature (1987-2010) that contains a figure representing pathology after diffuse or mixed pathology models of brain injury. At least 46 published manuscripts over this time period have shown histological assessment of pathology aligning with the temporal ridge following experimental TBI in the rodent.

| Year | Author | Title | Figure | Outcome Measure |
| --- | --- | --- | --- | --- |
| 1987 | McIntosh et al. | Traumatic brain injury in the rat: alterations in brain lactate and pH as characterized by <sup>1</sup>H and <sup>31</sup>P nuclear magnetic resonance | 2, 8 | Evan's Blue extravasation, Vulnerable brain region analysis |
| 1987 | McIntosh et al. | Traumatic brain injury in the rat: characterization of a midline fluid-percussion model | 6 | Subcortical hemorrhage |
| 1988 | McIntosh et al. | Magnesium deficiency exacerbates and pretreatment improves outcome following traumatic brain injury in rats: <sup>31</sup>P magnetic resonance spectroscopy and behavioral studies | 1 | Evan's Blue extravasation |
| 1989 | Cortez et al. | Experimental fluid percussion brain injury: vascular disruption and neuronal and glial alterations | 4 | Evan's Blue extravasation |
| 1989 | McIntosh et al. | Traumatic brain injury in the rat: characterization of a lateral fluid-percussion model | 8 | Evan's Blue extravasation |
| 1990 | McIntosh et al. | Effect of noncompetitive blockade of N-methyl-D-aspartate receptors on the neurochemical sequelae of experimental brain injury | 1 | Evan's Blue extravasation |
| 1991 | Hovda et al. | Diffuse prolonged depression of cerebral oxidative metabolism following concussive brain injury in the rat: a cytochrome oxidase histochemistry study | 1 | Cytochrome oxidase histochemistry |
| 1991 | Yoshino et al. | Dynamic changes in local cerebral glucose utilization following cerebral conclusion in rats: evidence of a hyper- and subsequent hypometabolic state | 1 | 2-deoxyglucose for glucose metabolic rate |
| 1992 | Hovda et al. | Secondary injury and acidosis | 5 | 2-deoxyglucose for glucose metabolic rate |
| 1992 | Soares et al. | Development of prolonged focal cerebral edema and regional cation changes following experimental brain injury in the rat | 1 | Vulnerable brain region analysis |
| 1993 | Hicks et al. | Mild experimental brain injury in the rat induces cognitive deficits associated with regional neuronal loss in the hippocampus | 2 | IgG extravasation |
| 1993 | Schmidt et al. | Regional patterns of blood-brain barrier breakdown following central and lateral fluid percussion injury in rodents | 6 | Biotinylated dextran amine for BBB breakdown |
| 1993 | Toulmond et al. | Biochemical and histological alterations induced by fluid percussion brain injury in the rat | 6 | Benzodiazepine binding for a neuronal marker |
| 1993 | Toulmond et al. | Prevention by eliprodil (SL 82.0715) of traumatic brain damage in the rat. Existence of a large (18 h) therapeutic window | 1 | Hematoxylin & Eosin |
| 1994 | Dietrich et al. | Widespread metabolic depression and reduced somatosensory circuit activation following traumatic brain injury in rats | 2 | 2-deoxyglucose for glucose metabolic rate |
| 1995 | Delahunty et al. | Differential consequences of lateral and central fluid percussion brain injury on receptor coupling in rat hippocampus | 4, 5, 6 | Cresyl violet |
| 1995 | Hicks et al. | Temporal response and effects of excitatory amino acid antagonism on microtubule-associated protein 2 immunoreactivity following experimental brain injury in rats | 3 | Microtubule associated protein immunohistochemistry |
| 1995 | Rink et al. | Evidence of apoptotic cell death after experimental traumatic brain injury in the rat | 2 | TUNEL+ stain |
| 1995 | Soares et al. | Inflammatory leukocytic recruitment and diffuse neuronal degeneration are separate pathological processes resulting from traumatic brain injury | 2 | Cresyl violet |
| 1995 | Soares et al. | Fetal hippocampal transplants attenuate CA3 pyramidal cell death resulting from fluid percussion brain injury in the rat | 2 | Cresyl violet |
| 1996 | Hicks et al. | Temporal and spatial characterization of neuronal injury following lateral fluid-percussion brain injury in the rat | 1 | Acid fuchsin, silver stain |
| 1996 | Saatman et al. | Prolonged calpain-mediated spectrin breakdown occurs regionally following experimental brain injury in the rat | 2 | Calpain-mediated spectrin breakdown immunohistochemistry |
| 1997 | Bareyre et al. | Time course of cerebral edema after traumatic brain injury in rats: effects of riluzole and mannitol | 1 | Vulnerable brain region analysis |
| 1997 | Iwamoto et al. | Investigation of morphological change of lateral and midline fluid percussion injury in rats, using magnetic resonance imaging | 1 | Magnetic resonance imaging |
| 1997 | Perri et al. | Metabolic quantification of lesion volume following experimental traumatic brain injury in the rat | 1 | TTC for succinate dehydrogenase activity |
| 1997 | Smith et al. | Progressive atrophy and neuron death for one year following brain trauma in the rat | 1 | Cresyl violet |
| 1998 | Conti et al. | Experimental brain injury induces regionally distinct apoptosis during the acute and delayed post-traumatic period | 2 | TUNEL+ stain |
| 1998 | Hulsebosch et al. | Traumatic brain injury in rats results in increased expression of Gap-43 that correlates with behavioral recovery | 2 | Growth-associated protein 43 immunohistochemistry |
| 1998 | Murakami et al. | Experimental brain injury induces expression of amyloid precursor protein, which may be related to neuronal loss in the hippocampus | 4 | Hematoxylin & Eosin |
| 1998 | Pierce et al. | Enduring cognitive, neurobehavioral and histopathological changes persist for up to one year following severe experimental brain injury in rats | 5 | Cresyl violet |
| 1999 | Di et al. | Fluid percussion brain injury exacerbates glutamate-induced focal damage in the rat | 1 | Hematoxylin & Eosin |
| 1999 | Hill-Felberg et al. | Concurrent loss and proliferation of astrocytes following lateral fluid percussion brain injury in the adult rat | 4 | Proliferating cell nuclear antigen positive cells for astrocytes, Glial fibrillary acidic protein for astrocytes |
| 2000 | Matsushita et al. | Real-time monitoring of glutamate following fluid percussion brain injury with hypoxia in the rat | 4 | TTC for succinate dehydrogenase activity |
| 2000 | Passineau et al. | Chronic metabolic sequelae of traumatic brain injury: prolonged suppression of somatosensory activation | 1, 2 | 2-deoxyglucose for glucose metabolic rate |
| 2001 | Harris et al. | Traumatic brain injury-induced changes in gene expression and functional activity of mitochondrial cytochrome c oxidase | 5 | Cytochrome oxidase enzyme histochemistry |
| 2001 | Sato et al. | Neuronal injury and loss after traumatic brain injury: time course and regional variability | 1 | Vulnerable brain region analysis |
| 2001 | Vink et al. | Small shifts in craniotomy position in the lateral fluid percussion injury model are associated with differential lesion development | 1 | Magnetic resonance imaging |
| 2002 | Bramlett et al. | Quantitative structural changes in white and gray matter 1 year following traumatic brain injury in rats | 2 | Hematoxylin & Eosin |
| 2004 | Cernak et al. | The pathobiology of moderate diffuse traumatic brain injury as identified using a new experimental model of injury in rats | 7 | Hematoxylin & Eosin |
| 2004 | Hallam et al. | Comparison of behavioral deficits and acute neuronal degeneration in rat lateral fluid percussion and weight-drop brain injury models | 3 | FluoroJade staining |
| 2004 | Singleton et al. | Identification and characterization of heterogeneous neuronal injury and death in regions of diffuse brain injury: evidence for multiple independent injury phenotypes | 5 | Fluorescent dextrans |
| 2005 | Schültke et al. | Neuroprotection following fluid percussion brain trauma: a pilot study using quercetin | 2 | Luxol fast blue-cresyl violet |
| 2005 | Van Putten et al. | Diffusion-weighted imaging of edema following traumatic brain injury in rats: effects of secondary hypoxia | 3 | Diffusion weighted imaging |
| 2009 | Lotocki et al. | Alterations in blood-brain barrier permeability to large and small molecules | 5 | Biotinylated dextran amine for BBB |
|  |  | and leukocyte accumulation after traumatic brain injury: effects of post-traumatic hypothermia |  | breakdown |
| 2010 | Hayward et al. | Association of chronic vascular changes with functional outcome after traumatic brain injury in rats | 2 | Cresyl violet, magnetic resonance imaging |
| 2010 | Yu et al. | Glial cell-mediated deterioration and repair of the nervous system after traumatic brain injury in a rat model as assessed by positron emission tomography | 1 | Positron emission tomography |

### Rat skull thickens at the temporal ridge

Naïve rat skulls were cleaned of all tissue using dermestid beetle larvae. Devoid of tissue, the prominence of the temporal ridges on the dorsal surface of the skull were evident (Figure 1). In fact, the overall shape of the external skull approximated a rectangular box, more so than the cylinder of the internal calvarium. Subsequent *in vivo* imaging was undertaken to demonstrate the relationship between the shapes of the skull and brain. Oblique sections of a 7T MRI in naïve rats were prepared to visualize the skull thickness with respect to the brain. Coronal (Figure 2A, 2B), horizontal (Figure 2C, 2D), and oblique sagittal (Figure 2E-2H) slices were all registered to pass through the temporal ridge of the skull (evident in black). Note that the thickness of the temporal ridge extended along the anterior to posterior length of the skull (Figure 2C). The internal surface of the bone underlying the temporal ridge remained continuous and uninterrupted with the rest of the skull, contacting the dura and brain as smooth surface devoid of macrostructure. To confirm the imaging results, coronal sections of rat skulls were taken from rostral to caudal and contour analysis was performed (Figure 3A-E). Measurements were taken along the temporal ridge and calvarium (*n*=4) of rat skulls (Figure 3F). The temporal ridge was found to be 75% thicker than the calvarium (t=4.36; p<0.01). Similar results occurred in the mouse, where skull thickness was greater at the temporal ridge than the sagittal bone (Supplementary Figure 1).

**Figure 1.**
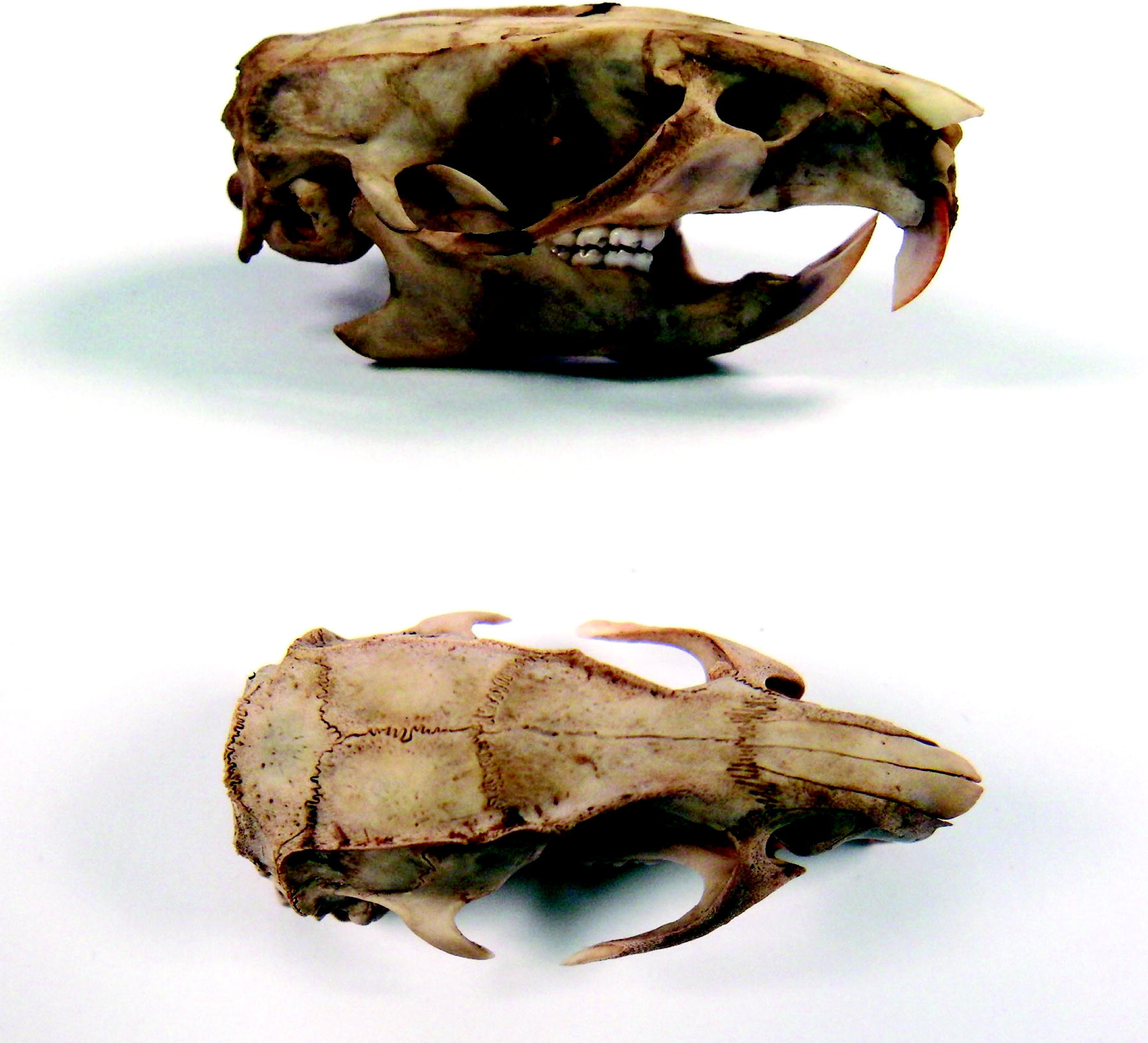
Rat heads were cleaned by flesh-eating beetle larvae until the skull was clear of all carrion. By gaining an anatomical perspective of the adult rat skull, the temporal ridge clearly protrudes on either side, indicating the increased bone mass in this region.

**Figure 2.**
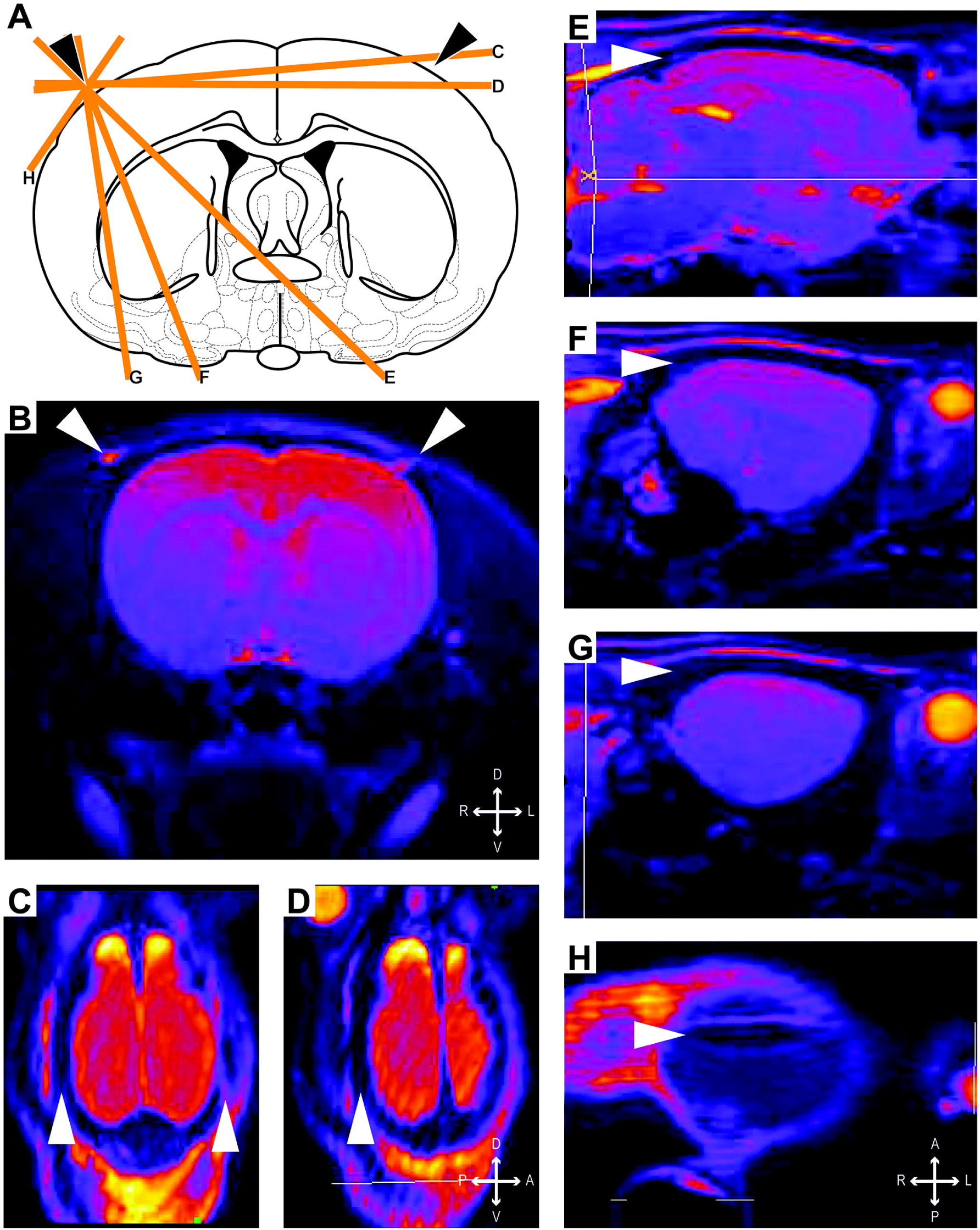
Oblique sections of naïve rat skull using a 7T MRI demonstrated the thickening of the rat skull along the temporal ridges. Coronal schematic (A) and section (B) present the conventional view of the rat brain. Oblique sections (C-H) are identified on the schematic at cross through the space beneath the temporal ridge. Temporal ridges are identified with the solid arrow head.

**Figure 3.**
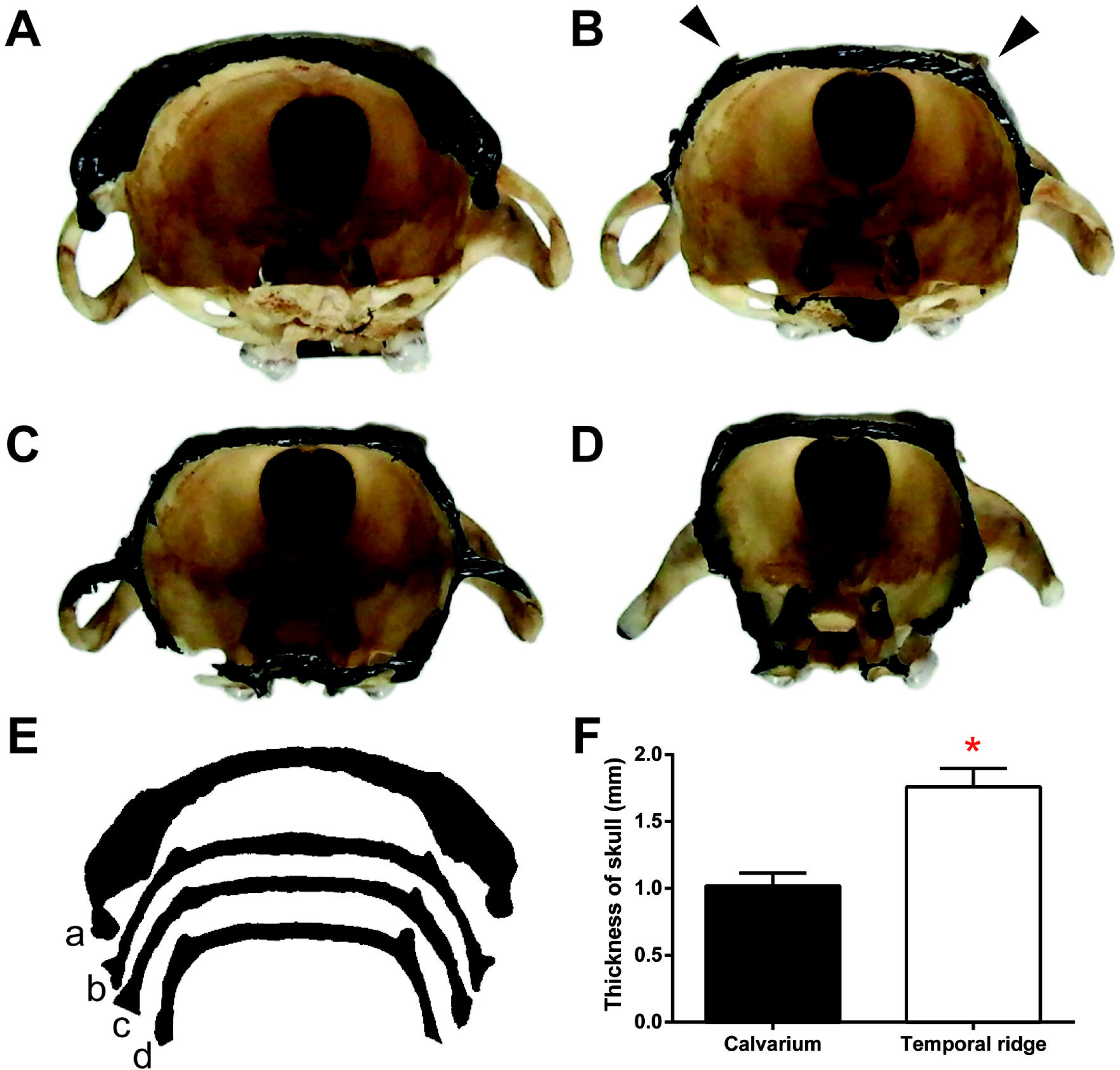
Local thickening of the temporal ridges is shown by contour analysis of coronal sections of the rat skull. (A-D) Photographs of the rat skull sections are enhanced by black ink on the sectioned edge. Solid arrowheads identify the temporal ridge. (E) Digitized outlines of the sectioned edges show the contour changes of the rat skull from posterior (a) to anterior (d). (F) Measurements were taken from coronal sections of rats to compare thickness of calvarium to the temporal ridge, where the temporal ridge was 75% thicker.

### FPI-induced argyrophilic neuropathology under the temporal ridge

Rats were diffuse brain-injured by mFPI and then survived to either 1, 2, 7 or 28 DPI. Brains were then collected, sectioned rostral to caudal and stained with silver to identify regions of neuropathology that develop following diffuse brain injury. Darker (black) stained regions on sections identify hyper-intense deposition of argyrophilic reaction product (de Olmos amino-cupric silver histochemical technique; Figure 4). For all time points post-injury, argyrophilia was evident lateral to midline, under the temporal ridge, and extended the rostral-caudal length of the brain. This length of pathology lies beneath the temporal ridge, with pathology evident bilaterally over the post-injury course. Specific brain regions (according to the Paxinos and Watson Rat Atlas) with argyrophilic staining were identified as the somatosensory cortex (S1BF), lateral portion of the hippocampus (CA3), and ventral posterior thalamus (Figure 5). This pattern of neuropathology occurred systematically across histological sections, with the deepest penetration of pathology in sections associated with the fluid pulse (center of the craniectomy). In some sections, areas of increased argyrophilic reaction product were inconsistent across the cortex (Figure 5), suggesting that a variable other than brain tissue properties may influence the pattern of pathology. The section presented for 1 DPI depicts a core devoid of argyrophilic staining, despite a penumbra of neuropathology. Yet, across the majority of histological sections, for all time points, tissue beneath the temporal ridge was preferentially vulnerable to neuropathology following mFPI. Similar neuropathology along rostral-caudal length of the brain under the temporal ridge occurred in the mouse (Supplementary Figure 2). Note that neuropathology may be diffuse and lateralized between hemispheres along the rostral-caudal length of the brain in both rats and mice (Figure 4, 5, Supplementary Figure 2).

**Figure 4.**
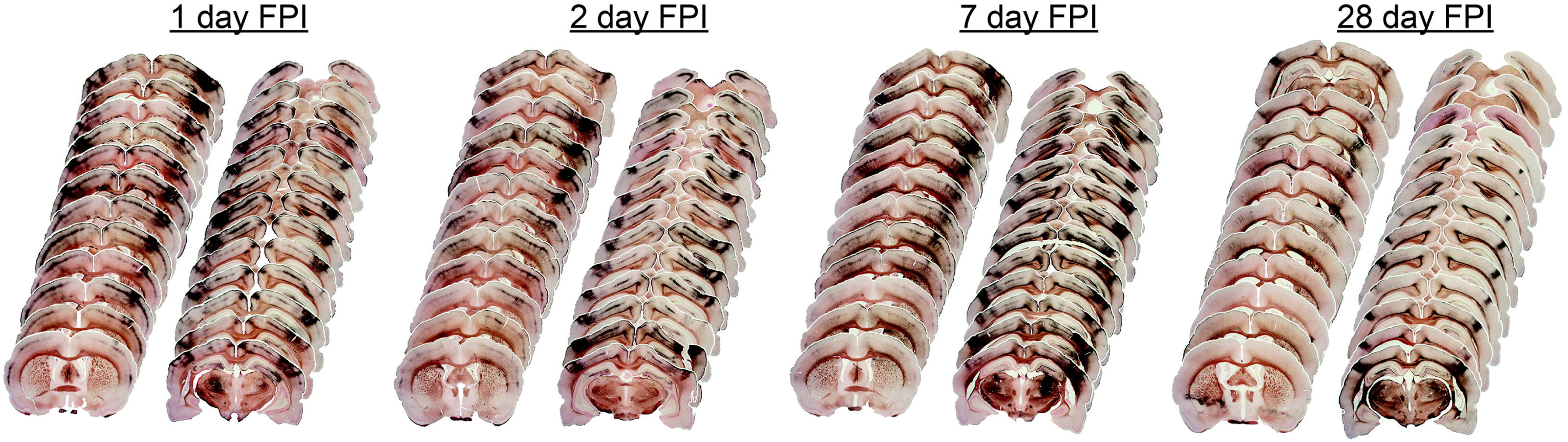
Histological sections of diffuse brain-injured rats are aligned rostral to caudal at 1, 2, 7, 28 days after midline fluid percussion injury (FPI). Neuropathology was identified by hyper-intense deposition of argyrophilic reaction product (amino-cupric silver histochemical technique; black) and occurred primarily along the rostral to caudal extent of sensorimotor cortex. Neuropathology appears to accumulate over 7 days post-injury and mostly subside by 28 days post-injury. Neuropathology may be diffuse and inconsistent between hemispheres along the rostral-caudal extent of the brain.

**Figure 5.**
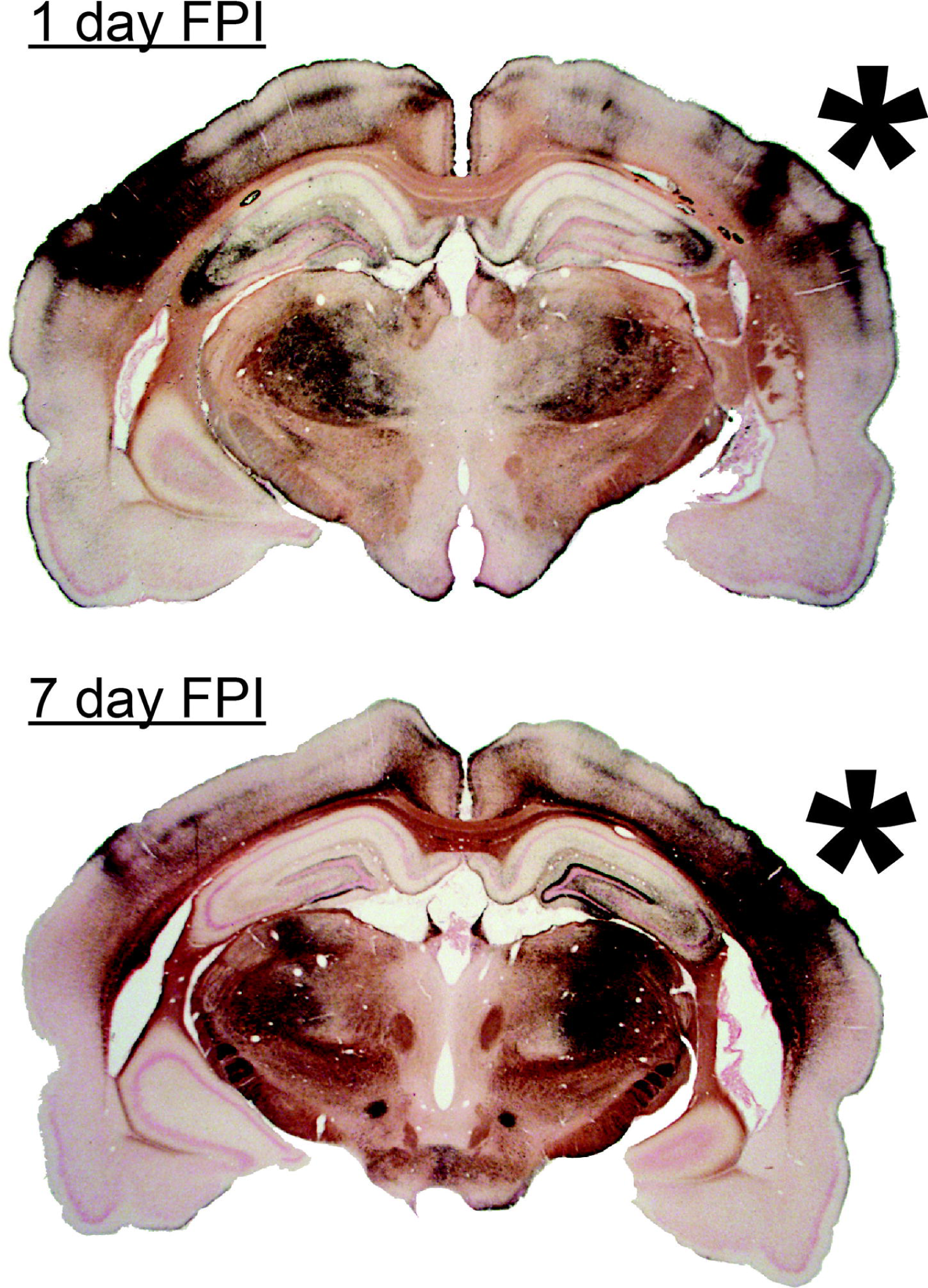
Tissue aligned with the temporal ridge (asterisk) showed increased deposition of argyrophilic reaction product (black) over time post-injury. The mechanical forces of diffuse brain injury reflect off the ventral skull into the temporal ridge, thereby inducing pathology in the lateral nuclei of the thalamus and lateral aspects of the hippocampus. Some sections (right side 1 day FPI) showed tissue spared of argyrophilic reaction at the temporal ridge, suggesting a non-neural variable may influence the pattern of pathology.

### Proposed biomechanical mechanism of rodent TBI

Consistent neuropathology occurred beneath and along the temporal ridge following mFPI, as represented in three dozen publications. Here, we propose a mechanism of injury induced by the mechanical forces of mFPI. Illustrated on a modified coronal MRI section (Figure 6A), mFPI is initiated (blue arrow) by the fluid pulse and pneumatic forces from the pendulum impact on the plunger of the fluid filled cylinder. This fluid pressure pulse, lasting only milliseconds, then produces mechanical force vectors that propagate throughout the brain (green arcs). As the wave propagates through the brain, the force vectors would reflect off the ventral skull, without causing damage. Reflected forces travel dorsal and lateral throughout the cranium, possibly towards the differential thickness of the skull at the temporal ridges (purple arcs). The differential thickness of the skull at the temporal ridges may act as either a pressure sink or a pressure barrier, which ultimately focuses injury-inducing forces back onto the tissue under the temporal ridge. This acts as a ‘pinch-point’ for vulnerable tissue and thereby contributes to the neuropathology observed acutely in the superficial cortical layers (red arrows). An alternative mechanism of injury would focus extracranial force vectors into the cranium through the temporal ridge, whereby the predicted pathology would initiate at the superficial cortical layers and diffuse ventrally from those points. For either proposed mechanism, the applied forces may remain localized to the originating cerebral hemisphere(s), such that lFPI is lateralized compared to mFPI. We favor the intracranial mechanics proposed mechanism since the pathology shows the largest arc across the cortex at superficial layers.

**Figure 6.**
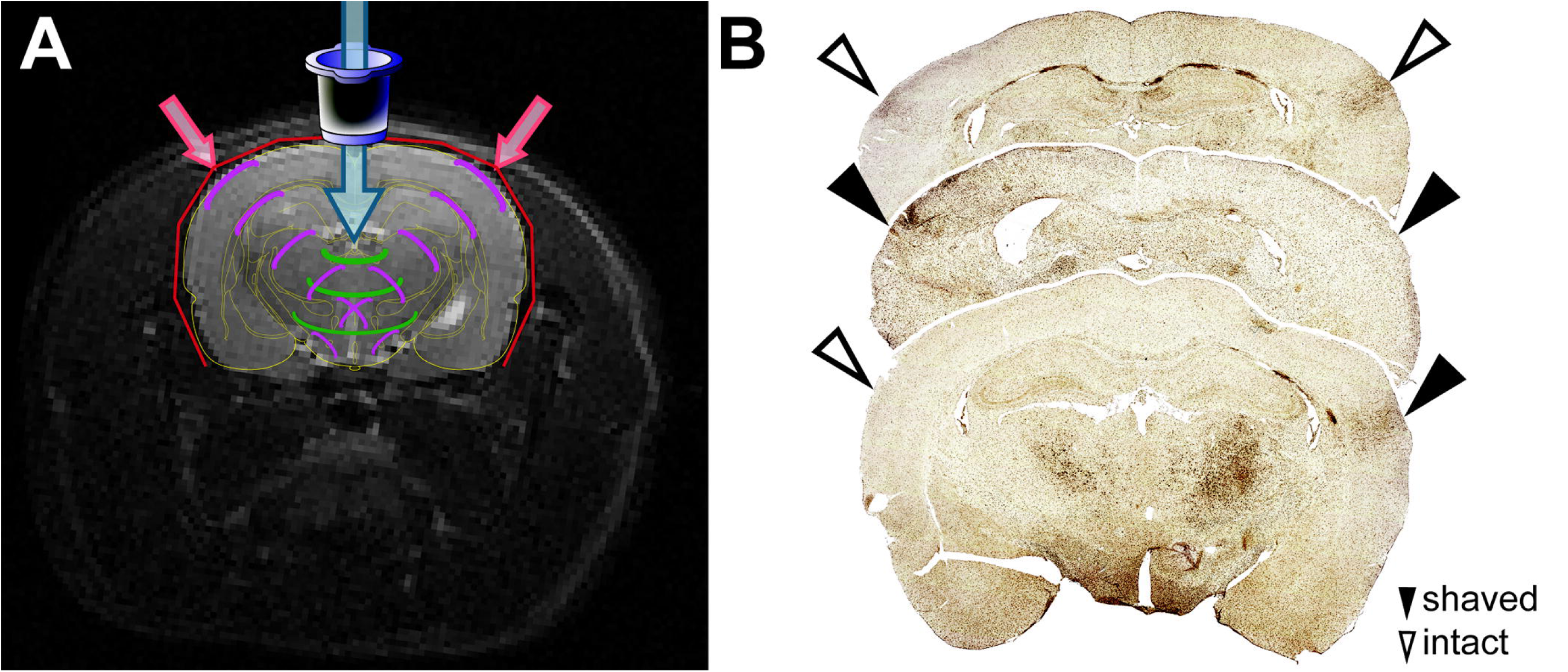
Proposed biomechanical mechanism of rodent TBI. (A) Schematic of mechanical forces following midline fluid percussion injury (mFPI) that induce neuropathology. The fluid pulse (blue arrow), generated from the impact of the pendulum on the plunger of a fluid-filled cylinder lasts only milliseconds, and travels through the injury hub into the extradural space producing mechanical forces that propagate throughout the brain (green arcs). Mechanical forces are then reflected off the ventral portions of the skull, and travel throughout the cranium back to the dorsal and lateral portions of the skull (purple arcs). Upon reaching the cortex, increased thickness of the temporal ridge provides a pressure sink or pressure barrier, which ultimately focuses injury-inducing forces on the tissue beneath the temporal ridge, acting as a ‘pinch point’ and resulting in observed neuropathology (red arrows). (B) To support this proposed mechanism, temporal ridges were shaved unilaterally or bilaterally to approximate the thickness of the calvarium prior to injury. Rats then received mFPI and were prepared for Iba-1 immunohistochemistry at 7 DPI to identify areas of neuroinflammation. Rats who received no shaving to the temporal ridge (top brain slice) show focal increase of microglial activation. However, when the temporal ridge was unilaterally (bottom brain slice) or bilaterally (middle brain slice) shaved, an absence of focal neuroinflammation corresponded with the shaved hemisphere(s). Open arrow heads indicate hemispheres with an intact temporal ridge; solid arrow heads indicate hemispheres with a shaved temporal ridge.

To support this proposed mechanism, one or two temporal ridge(s) were shaved down prior to mFPI to approximate the thickness of the calvarium. Rats then received moderate mFPI and were prepared for immunohistochemistry at 7 DPI. Brains were collected, sectioned and immunostained with Iba-1 to identify concentrated areas of neuroinflammation, indicative of ongoing neuropathology [61, 63]. Representative histological sections show (Figure 6B) increased focal neuroinflammation (Iba1+ microglial activation) corresponding to hemispheres with an intact temporal ridge. Rats with a single temporal ridge intact showed an absence of focal neuroinflammation in the hemisphere corresponding with the side of the shaved temporal ridge (Figure 6B, bottom brain section). Rats with both temporal ridges intact showed the predicted concentrated microglial activation in cortical areas beneath the temporal ridge (Figure 6B, top brain section). Rats with both of the temporal ridges shaved showed diffuse microglial activation (Figure 6B, middle brain section).

## Discussion

Here we present evidence to explain the repeatedly documented pathology that occurs lateral to the injury induction site in experimental TBI. In the literature, dozens of articles contain figures showing aspects of neuropathology preferentially localized to tissue beneath the temporal ridge following mFPI and other models of closed head injury. Having observed the location of this curious pathology in the literature and our own work, we characterized the increased thickness of the skull at the temporal ridge and the skull:brain relationship using MRI, skulls, and contour imaging of coronal sections from naïve rat skulls. Furthermore, we show that following mFPI, neuropathology identified with silver stain occurs in cortical tissue running the rostral-caudal length of the temporal ridge, and includes the S1BF, hippocampus (CA3) and ventral thalamus in the dorsal-ventral projections of mechanical force. We thus propose a new biomechanical mechanism of experimental TBI mechanical forces to explain this predictable and consistent neuropathology. To support this proposed mechanism, we unilaterally or bilaterally shaved the temporal ridge prior to mFPI and demonstrate the absence of focal neuroinflammation under the single shaved ridge. Together, these data provide new insight into the mechanical contribution of the rodent skull, rather than specific tissue properties, in the context of experimental brain injury.

Since the implementation of FPI, multiple groups have hypothesized the biomechanics of the injury. However, in each of these reports, injury induction parameters were the primary considerations, rather than the resultant neuropathology [14, 64]. As shown in Figure 3, the rat skull is not uniform in thickness and could thus influence mechanical force trajectories applied to the head and skull. As Dixon *et al*. noted, “*fluid moves through the epidural space of the brain after FPI*” [5]. Lighthall *et al*. further explained how the complex pattern of tissue deformation occurs (hypothesized to be related to gray:white matter interfaces), thus, causing diffuse neuropathology [64]. For this model, it appears that the applied mechanical force increased strain and stress on neuronal tissue, with the likely consequence of tissue forced against different parts of the skull. Recent *in silico* modeling of lFPI included a 3-layer hexahedral, element-based skull module with varying Young modulus for each layer [65]. The uniformly thick skull appeared not to influence the results, which showed local forces at the site of injury and secondarily at the ventral aspect of the brain, as proposed here. An opportunity exists for future finite element models to consider differential skull thickness, which may confirm or refute the result of reflected vector forces, as indicated by the neuropathology. While each of these studies describes how the fluid pulse interacts with and thus deforms neuronal tissue, they do not explain the unique pattern of neuropathology observed post-injury. Moreover, these studies do not take into consideration the forces as they reflect off the skull and travel through neuronal tissue. No considerations are given to the differential skull thickness or its shape. As we characterize here, the rat skull is not a uniform thickness and therefore would not absorb or reflect forces equally. The temporal ridge protrudes externally along the rostral-caudal skull axis, potentially increasing bone rigidity. The calvarium in contact with the dura and brain is contoured, smooth, and free of any protrusions. One hypothesis to explain the temporal ridge relates to the attachment and early use of muscles of mastication [66-69]. Suckling and eating following birth can increase strains along the rostral-caudal axis of the rodent skull sufficiently to influence the development of the temporal ridge. In adulthood, the temporal ridge is 75% thicker than the rest of the calvarium, thus providing stiffness along this point of the skull.

Evidence across the literature indicates that tissue directly underneath the temporal ridge, rather than the impact site, routinely shows TBI-induced neuropathology. We show evidence of this pathology throughout the rostral-caudal extent of coronal sections. This neuropathology, evidenced in multiple histological outcomes by the Neurotrauma community, is multifocal and preferentially in brain regions within proximity to the temporal ridge [6, 11, 18, 24, 37, 40, 56, 70-108]. It is essential to note that the studies listed in Table 1, in corroboration with the silver-stained histological sections in this communication, represent multiple Neurotrauma laboratories across the world, over decades, and are the product of numerous surgeons. Thus, the representative neuropathology associated with the temporal ridge is more likely a feature of injury forces in the rodent brain, rather than spurious surgical variation. We contend that the curious neuropathology results from brain tissue forced into the temporal ridge, as the injury forces reflect off the ventral skull. Transmission through ventral structures and reflection off the ventral skull would predict optic nerve damage, as reported by the Povlishock group in mFPI [109, 110]. In some cases, identified in Figure 5, the neuropathology is not uniform along the cortex near the temporal ridge, suggesting that tissue may warp non-uniformly at the temporal ridge of the skull, thereby sparing some cortical tissue from the full forces of injury. Aspects of mechanical forces that may damage the brainstem, as has been shown for FPI in the cat, may exist and need further investigation [111, 112]. Further, gyrencephalic brains may absorb or reflect forces in a manner that minimizes the influence of an overall thicker skull. In this communication, we argue that the relationship of the temporal ridge to the injured tissue is not coincidental or due to the differences in surgical technique.

For brain injury models that intentionally penetrate the dura, such as controlled cortical impact (CCI), neuropathology is expected and localized primarily to the impact site. In these focal injury models, cavitation occurs at the injury site, with a penumbra of tissue damage [113]. However, neuropathology after focal CCI spreads into diffuse pathology of the contralateral hemisphere, with accumulation of argyrophilia in cortical areas under the temporal ridge by 7 DPI [113]. Alternatively, if the craniotomy for FPI is performed over the temporal ridge, then no overt pathology is observed [114, 115]. Different positions of the craniotomy on the skull, or non-perpendicular injury hub angles, may unequally distribute injury forces between the two hemispheres. At the present time, non-contact brain injury models, such as blast injury, have not reported substantial cortical neuropathology, yet it is predicted to be localized in regions tracking to the temporal ridge. Finally, cortical neuropathology in murine diffuse TBI models also traversed the temporal ridge, as presented for mFPI (Supplementary information), further supporting a role for the temporal ridge in the proceeding neuropathology.

Closed head injury in the rodent results in neuropathology preferentially in the somatosensory cortex (S1BF), hippocampus (CA3), and ventral thalamus. These three regions lie in a wedge or arc from the ventral midline aspect of the skull base towards the temporal ridge. For our biomechanical model of brain injury forces, these regions may not be uniquely vulnerable to TBI pathophysiology, but rather casualties along the force vectors (Figure 6A). By removing one temporal ridge, the neuropathology becomes localized to a single hemisphere, whereas removing both temporal ridges may retain the bilateral pathology due to the geometric shape of the skull, regardless of the temporal ridges. Further, the neuropathology in these regions would predict neurological and behavioral impairments, including somatosensory and cognitive impairment. With regard to the S1BF, whisker sensitivity and somatosensory dysfunction have been reported [29-34, 116, 117]. Further, these neurological impairments coincide with altered circuitry that involves both the S1BF and ventral thalamus. Cognitive performance involving short, long, and working memory, using multiple established mazes, is impaired following FPI [10, 32, 35-45]. These cognitive impairments may involve CA3 processing to perform object pattern completion, cue retrieval in fear conditioning, episodic memory, and spatial memory [12, 118-125]. Thus, the neuropathology associated with the temporal ridge to include the S1BF, CA3, and ventral thalamus manifests with impairments in behavioral performance post-injury.

The pathology and neurological impairments after diffuse TBI are associated with disrupted circuitry. Our ongoing work uses experimental models, such as rodent FPI, to define injury and repair mechanisms. In this communication, we show one, amongst many possible, explanations for the unique pathology observed in select brain regions of the diffuse injured rodent brain. Intracranial force measurements in multiple locations using physical probes would compromise the calvarium and fundamentally change the injury, which further restricts the ability for biomechanical modeling. Additionally, we are unaware of an anatomically accurate finite element model of the rat skull that would permit rapid evaluation of brain injury biomechanics; future endeavors can develop and incorporate skull anatomy into finite element models. Overall, the interpretation of the injury forces causing the observed neuropathology need not be mutually exclusive. Regional heterogeneity in architecture, cell composition, proximity to white matter, biochemistry (glutamate receptors), and enzymatic profiles contribute to the observed pathology. Further, the observed pathology cannot be fully explained by the time course of a single variable (e.g. blood pressure, herniation, excitotoxicity, potassium level). The unique structure of the rodent skull may converge these alterations on specific brain regions. Alternatively, tissue properties that impart vulnerability to injury may propagate force vectors, but the pathology under the temporal ridge adds circumspect to this conjecture. The temporal ridge is a likely contributor to the genesis of diffuse brain injury pathology and neuroinflammation, which may be unique to the rodent.

By analyzing the curious pathology of mFPI and relating the biomechanical model to other closed head injury models, laboratory studies can take advantage of localized pathology, without overt cavitation, to explore post-traumatic reorganization and repair of the cortex, hippocampus, and ventral thalamus [61, 62]. By focusing on neuronal circuits that regulate somatosensory and cognitive function, investigators may continue to advance our understanding of the disease process that dismantles, repairs, and regenerates circuits in the brain. These results encourage increased precision in defining regions of injury, where cortex analysis insufficiently addresses the regional heterogeneity. It remains likely that gene expression, synapse loss, receptor expression, microglial activation, and cell death would vary within the cortex, as demonstrated for microglia deramification [126]. The pathology of diffuse TBI is the summation of (1) the mechanical forces of the primary injury, (2) the biomechanics of the impacted substrates (e.g. temporal ridge, tissue modulus of the brain), (3) the subsequent signaling cascades, and (4) secondary injuries (e.g. hypoxia, seizures). Ultimately, the acute events of TBI initiate a disease process that leaves individuals with debilitating symptoms, which impair their quality of life [2]. Through continued investigation of the cellular, molecular, structural, functional, and behavioral consequences of TBI, we strive to improve the quality of life for our patients by advancing diagnostic techniques and therapeutic interventions.

## Supporting information

Supplemental Methods, Results, Discussion and Figure Legends

## Acknowledgements

We are grateful to Amanda Lisembee, Kelley Hall, Mark Abromovitz, Sarah Ogle, Daniel R. Griffiths, Charlotte Denman, and Anna Learoyd for technical and surgical expertise. Further, these studies depended on the carrion beetles supplied by James J. Krupa, Ph.D., Department of Biology, University of Kentucky. Additionally, we thank Rory Young for his contributions to the publications analysis. Finally, we would like to acknowledge Bruce G. Lyeth for support and intellectual discussion that shaped the direction of this paper.

## Human and Animal Rights

All work included in this study was conducted in compliance with the applicable institutional ethical guidelines for the care, welfare, and use of animals.

## Data Availability

The data supporting the findings of this study are available from the corresponding author upon reasonable request.

