## Supplemental Methods, Results, Discussion and Figure Legends for "Spatial distribution of neuropathology and neuroinflammation elucidate the biomechanics of fluid percussion injury"

### Supplementary Methods

Male C57BL/6 mice (Jackson Laboratories, Inc., Bar Harbor, ME) were used for all experiments. The animals were housed in a 12 h light/12 h dark cycle at a constant temperature with food and water available *ad libitum* according to the Association for Assessment and Accreditation of Laboratory Animal Care International. After surgery (described below) animals were evaluated daily for postoperative care by a physical examination and documentation of each animal's condition. Animal care was approved by the Institutional Animal Care and Use Committees at Virginia Commonwealth University (Richmond, Virginia).

Mice (20-26 g) were subjected to mFPI consistent with methods previously described [59]. After anesthesia induction, each animal's thigh was shaved for intra-operative physiological monitoring and placed in a stereotactic frame (David Kopf Instruments, Tujunga, CA) fitted with a nose cone to maintain anesthesia with 1–2% isoflurane in 100% $O_2$. A thermostatically controlled heating pad (Harvard Apparatus, Holliston, MA) was then placed under the animal and set to monitor the rectal temperature and, via feedback control, maintain the body temperature at 37°C during the surgery. Pulse rate, respiratory rate and blood oxygenation were monitored intra-operatively via pulse oximetry using a thigh sensor (STARR Life Sciences Corp.; Oakmont, PA) to ensure the maintenance of normal physiologic homeostasis with the exclusion of animals sustaining any physiological anomaly.

A midline sagittal incision was made to expose the skull from bregma to lambda. The skull was cleaned and dried and a 3.0 mm circular craniotomy was then made along the sagittal suture midway between bregma and lambda, leaving the underlying dura intact. A sterile Leur-Loc syringe hub was then cut away from a 20-gauge needle and affixed to the craniotomy site using cyanoacrylate. Upon confirming the integrity of the seal between the hub and the skull, dental acrylic was then applied around the hub to provide stability during the induction of injury. After the dental acrylic hardened, the scalp was sutured around the hub, topical bacitracin and lidocaine ointment were applied to the incision site, and the animal was removed from anesthesia and monitored in a warmed cage until fully ambulatory (approximately 60–90 minutes).

For the induction of injury, each animal was re-anesthetized with 4% isoflurane in 100% $O_2$, and the male end of a spacing tube was inserted into the hub. The female end of the hub-spacer assembly, filled with normal saline, was attached on to the male end of the fluid percussion apparatus (Custom Design & Fabrication; Virginia Commonwealth University; Richmond, VA). An injury of mild to moderate severity (1.7±0.04 atmospheres) was administered by releasing a pendulum onto a fluid-filled piston to induce a brief fluid pressure pulse upon the intact dura. The pressure pulse measured by the transducer was displayed on a storage oscilloscope (Tektronix 5111, Beaverton, OR), and the peak pressure was recorded. After injury, the animals were visually monitored for recovery of spontaneous respiration. The hub and dental acrylic were removed *en bloc*, and the incision was rapidly sutured before recovery from anesthesia/loss of responsiveness. Topical bacitracin and lidocaine were then applied to the closed scalp incision. The duration of transient loss of responsiveness was determined by measuring the time it took each animal to right itself from a supine position. After recovery of the righting reflex, animals were placed in a warmed holding cage to ensure the maintenance of normothermia and monitored during recovery before being returned to the vivarium.

At the pre-determined end points, mice were intraperitoneally injected with an overdose of sodium pentobarbital and then transcardially perfused with heparinized normal saline followed by 4% paraformaldehyde. Following decapitation, the heads were stored in a fixative solution containing 15% sucrose for 24 h, after which the brains were removed, placed in fresh fixative, and shipped for histological processing to Neuroscience Associates Inc. (Knoxville, TN). The mouse brains were embedded into a single gelatin block (Multiblock® Technology; Neuroscience Associates). Individual cryosections containing all the mouse brains were mounted and stained with the de Olmos aminocupric silver technique according to proprietary protocols (Neuroscience Associates) to reveal argyrophilic reaction product, which localized to neurons and neuronal processes, counterstained with Neutral Red, and then cover-slipped. The stained sections were digitized and returned to the laboratory. Every sixth section from the anterior commissure through the substantia nigra was masked from the background and overlaid on the remaining sections from the same brain.

#### Supplementary Results

Naïve mouse skulls were cleaned of all tissue using dermestid beetles. Devoid of tissue, the prominence of the temporal ridge along the dorsal surface was evident. Similar to that of the rat skull, the rectangular shape of the skull was identified in the naïve mouse skulls. To confirm homology between the thickening along the temporal ridge observed in rat and mouse skulls, coronal sections were taken, and contour analysis was conducted (Supplementary Figure 1A-B). Measurements were taken from rostral to caudal ($n$=2) of mouse skulls. The temporal ridge was found to be 43% thicker than the calvarium (Supplementary Figure 1C).

Mice received diffuse brain injury by mFPI and then aged to either 2 or 7 DPI. Brains were then collected, sectioned rostral to caudal and stained with silver stain to identify regions of neuropathology. Darker regions on sections presented in Supplementary Figure 2 represent deposition of argyrophilic reaction product (amino-cupric silver histochemical technique) and are preferentially located lateral to the injury site. Furthermore, the neuropathology extended under the rostral-caudal length of the temporal ridge, with the areas of the most intense argyrophilic staining at the location of injury impact.

#### Supplementary Discussion

Here we provide further evidence for lateralized neuropathology following mFPI that is not specific to the rat. Mice also showed a thickening of the skull at the temporal ridge (43% in mouse compared to 75% in rat) and an accumulation of argyrophilic staining along the sensorimotor cortex underlying the rostral-caudal extent of the brain. Additionally, this neuropathology coincides with acute motor deficits, similar to those observed in the rat. Motor deficits have been reported to mice subjected to mFPI using tests of neuromotor function such as the beam balance, beam walk, rotarod, and inclined plane test [12, 16, 20, 124]. Following mFPI in the mouse, these neurobehavioral deficits are most evident in the first few days post-injury, when compared to uninjured control animals. Mice subjected to mFPI have also been reported to have acute deficits in retrograde memory recall in the Morris water maze [123] and novel object recognition task [20], paralleling CA3 dependent processing impairments in brain-injured rats. Due to the similarities between rat and mouse, we propose that our mechanism for the biomechanics of FPI is applicable to both species, and perhaps to all species with a pronounced temporal ridge. Yet, the argyrophila at the impact site in mice may represent the consequences of thinner temporal ridge than in rats. Investigators may take advantage of localized pathology, without overt cavitation, to refine tissue, histological, and behavioral analyses.

#### Supplementary Figure 1

Local thickening of the skull at the temporal ridges is shown by contour analysis of coronal sections of the mouse skull (A-B). The temporal ridge was 43% thicker than the calvarium (C; t-test not significant).

#### Supplementary Figure 2

Histological sections of diffuse brain-injured mice are aligned rostral to caudal at 2 and 7 days after midline fluid percussion injury (FPI). Neuropathology was identified by hyper-intense deposition of argyrophilic reaction product (amino-cupric silver histochemical technique; black) and occurred primarily along the rostral to caudal extent of sensorimotor cortex. Neuropathology observed showed similarity to that of the diffuse injured rat brain, suggesting similar injury biomechanics between the two species.
