## Supplementary figures and images for "Spatial distribution of neuropathology and neuroinflammation elucidate the biomechanics of fluid percussion injury"

### Supplemental Figure 1

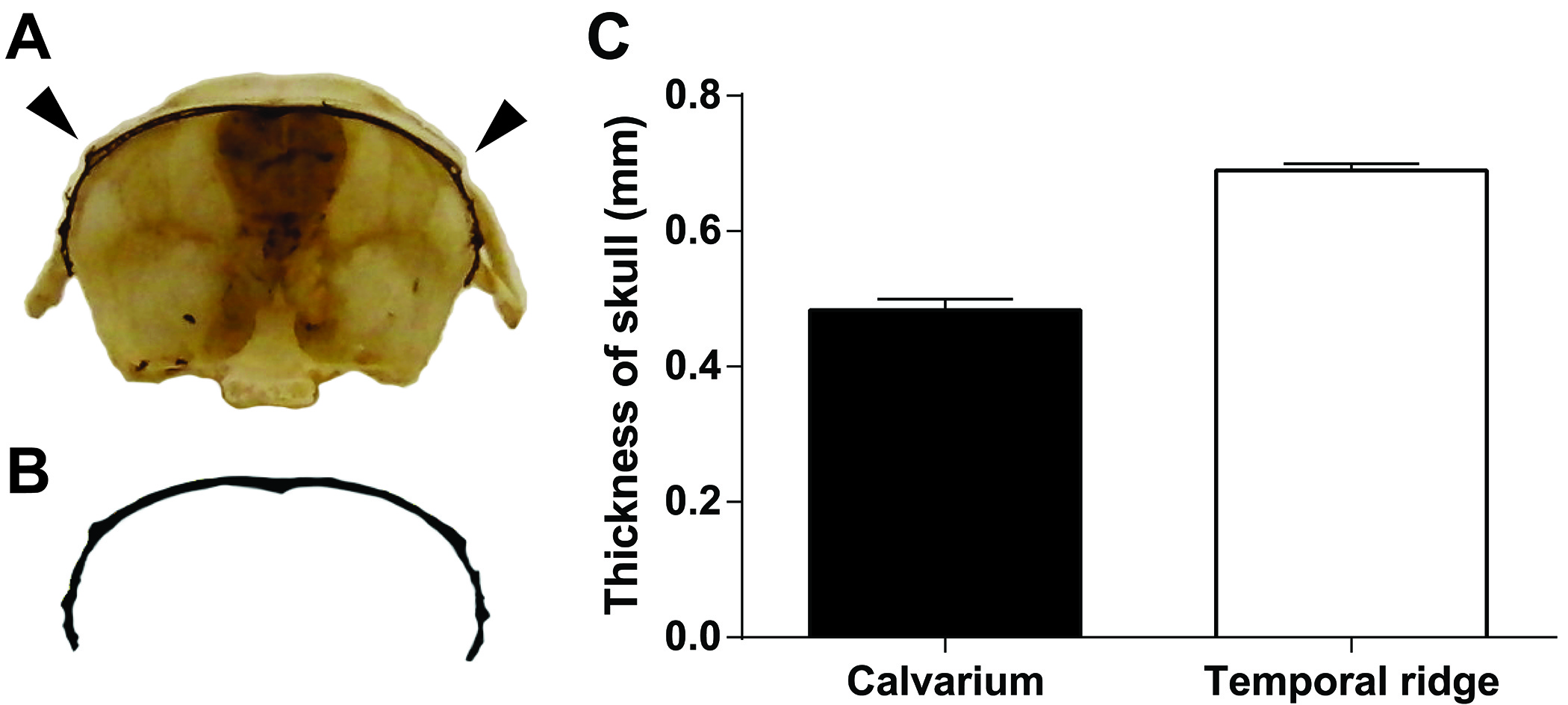

### Supplemental Figure 2

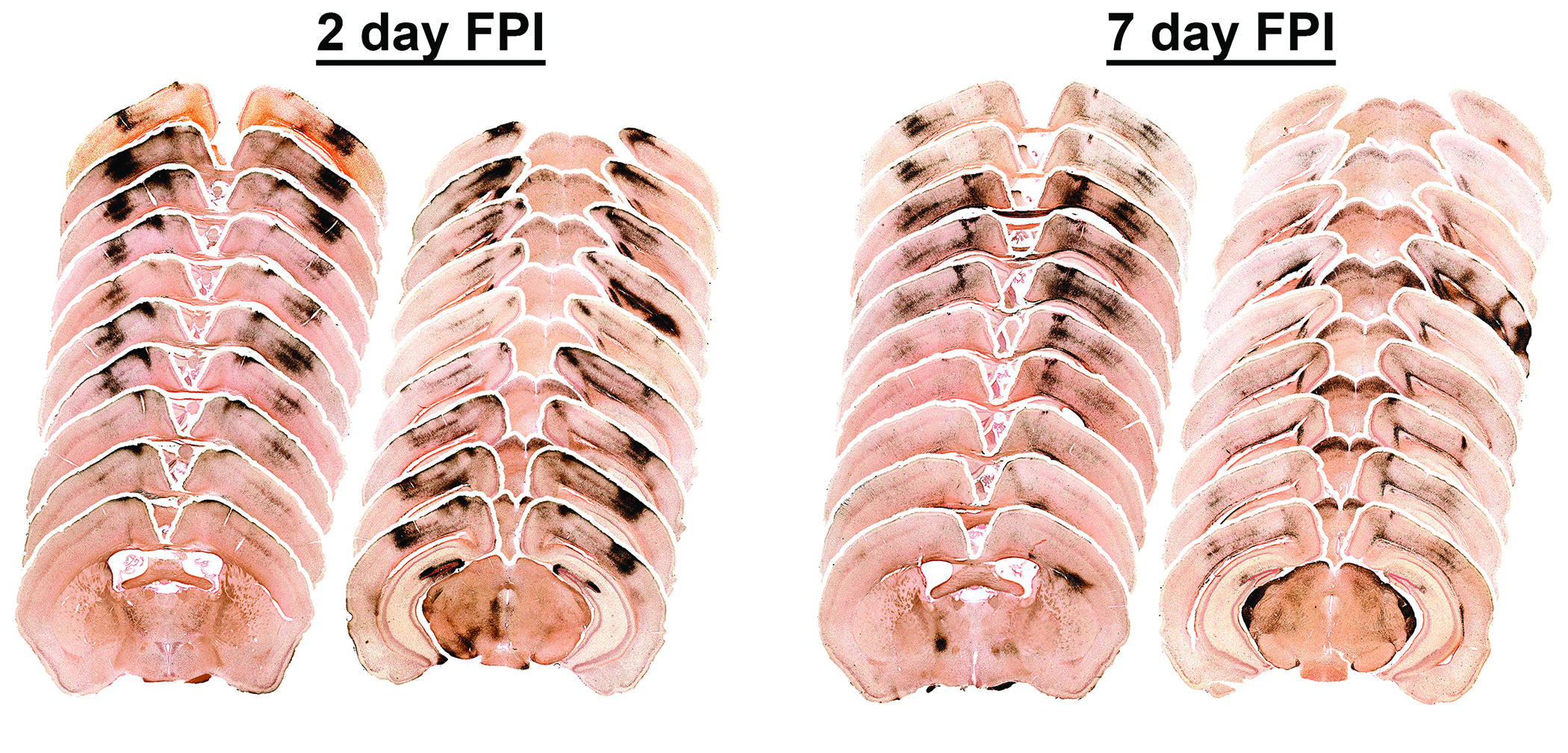
